# Mass Spectrometry based Metabolomics Deciphered Metabolic Reprogramming That Was Required for Biofilm formation in Uropathogenic *Escherichia coli*

**DOI:** 10.1101/672592

**Authors:** Haitao Lu, Yumei Que, Xia Wu, Tianbing Guan, Hao Guo

**Author notes:** The authors are equally contributed to this article (Haitao Lu, Yumei Que). Corresponding author., Haitao Lu Ph.D. Professor, Key Laboratory of Systems Biomedicine (Ministry of Education), Shanghai Center for Systems Biomedicine, Shanghai Jiao Tong, University, Shanghai 200240, China, Email.

## Abstract

Biofilm formation plays a key role in many bacteria causing infections, which mostly accounts for high-frequency infectious recurrence and antibiotics resistance. In this study, we sought to compare modified metabolism of biofilm and planktonic populations with UIT89, a predominant agent of urinary tract infection, by combining mass spectrometry based untargeted and targeted metabolomics methods, as well as cytological visualization, which enable us to identify the driven metabolites and associated metabolic pathways underlying biofilm formation. Surprisingly, our finding revealed distinct differences in both phenotypic morphology and metabolism between two patterns. Furthermore, we identified and characterized 38 differential metabolites and associated three metabolic pathways involving glycerolipid metabolism, amino acid metabolism and carbohydrate metabolism that were altered mostly during biofilm formation. This discovery in metabolic phenotyping permitted biofilm formation shall provide us a novel insight into the desperation of biofilm, which enable to develop novel biofilm based treatments against pathogen causing infections, with lower antibiotic resistance.

---

Bacterial biofilms are fundamentally structured in a manner of a self-produced matrix of extracellular polymeric substance (EPS) with the extracellular DNAs, proteins and polysaccharides (Hung et al., 2013). Biofilm formation drives the bacterial cells to persist survival and prolong their life-span, which has been observed to play a key role in many human infections with high-frequency antibiotic resistance (Costerton et al., 1995). Urinary Tract Infection (UTI) is a common, high-recurrence or even life-threaten infectious disease, which is mostly caused by uropathogenic *Escherichia coli* (UPEC). UPEC prefers to form biofilm and trigger the infections recurrently, thereby leading to uneasily eradicate UPEC strains by the treatment of antibiotics against UTI (Hung et al., 2013; Lv et al., 2014; Yan et al., 2015). Basically, biofilm formations of bacteria have distinct stages involving attachment, micro colony formation and maturation, which are almost agreeable among different bacterial species. However, it still remains unclear if small-molecule metabolites and associated metabolism were required for biofilm formation and degradation. Our previous work verified that modified metabolism was observed to closely associate with the virulence expression of UPEC strains so as to promote the progression of UTIs, herein, reprogrammed metabolism might be the key factor for biofilm formation in terms of such structure directly contributes to high-pathogenicity and recurrence of UPEC strains during causing infections. Pellicle is a classical growth-mode of biofilm *in in vitro,* which is formed at the air-liquid interface without anchoring themselves to any solid surfaces (Hung and Henderson, 2009; Hung et al., 2013). UTI89 strain is a clinical UPEC strain that was originally isolated from the bladder of a woman with UTI, this stain is capable of forming a floating pellicle-biofilm in YESCA medium. To better understand the molecular phenotype of biofilm formation triggered by UPEC strain from a metabolic perspective, we aimed at investigate the biofilm formation by integrating staining assay, cytological observation with metabolomics method (Figure 1A).

**Figure 1.**
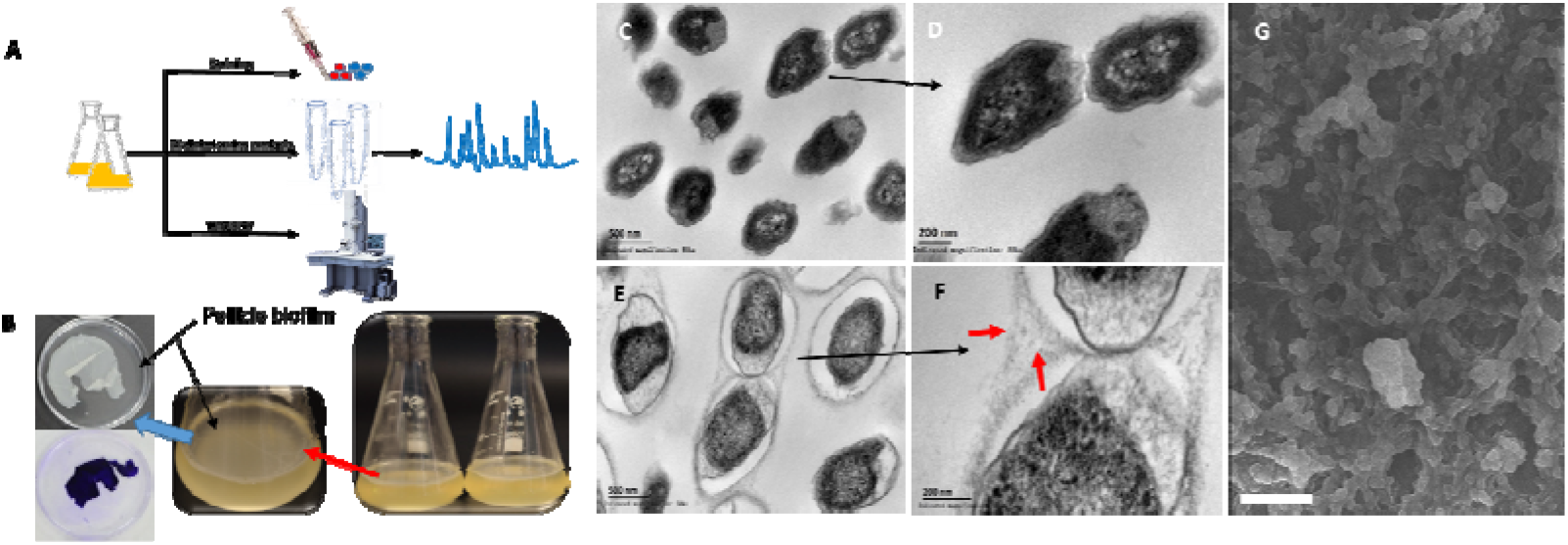
Combinational strategy characterized biofilm phenotype of UPEC UTI89 by integrating metabolomics method (A),. staining assay (B) with cytological observation (C-G).

## Materials and Methods

### Reagents, Bacterial Strains and culture process

UTI89 was firstly incubated with LB broth (Miller’s LB) (BD, Franklin Lakes, USA). The biofilm formation culture was facilitated statically in YESCA medium (1% Calamine Acids and 0.12% yeast extract) in 96-well plastic plates for Congo red and crystal violet for staining, as well as conical flask for metabolite enrichment and extraction. Briefly, one colony UTI89 strain was incubated with 7ml LB broth for 5h on a shaker, and then the bacterial cells were diluted 100-fold into fresh YESCA medium, then incubated at 30° for 96 h statistically to trigger biofilm formation. The Congo red concentration was diluted in YESCA medium at 50ug/ml. All the reference compounds for targeted metabolite assay were purchased from Sigma Company (Sigma, St. Louis, USA)

### Crystal Violet Staining Assay

Biofilm formation was quantified by crystal violet staining assay. Biofilm was initiated as previous described. The mature biofilm was washed with PBS for 3 times and then fixed biofilm with methanol for 15 minutes.

After the methanol volatilized, 100 ul 1% crystal violet was added to the biofilm for 20 min. The dye wells with crystal violet were washed with PBS for 5 times. Bound crystal violet was solubilized with 100ul 33% acetic acid for 30 min whilst shaking slowly. ODS value was measured via microplate reader at 570 nm. Statistical significance was determined by 2-tailed Student’s T-test with a threshold p-value <0.05

### CFU Assay

Each biofilm-sample was completely dispersed by manual-homogenizer in PBS for 20 times with the same physical pressure. The biofilm suspensions and planktonic cells were used to measure the CFU value. The following steps are recorded in reference^3^

### SEM assay

Mature biofilm and the planktonic cells were fixed with 2.5% glutaraldehyde for more than 3 hours. Next, they were washed for three times with PBS, and then the specimen was first to be dehydrated by gradient-concentration ethanol (30% 50% 70% 80% 90 % and 100%) for 5min at each step. After that, the specimen was dehydrated again by the gradient-concentration tertiary butanol (70% 90% 95% and 100%). At last, the dehydrated specimen was coated with gold-palladium and observed in SEM (HITACHI S-3400N).

### TEM assay

Mature biofilm and the planktonic cells were fixed with 2.5% glutaraldehyde for more than 3 hours. All the samples were embedded in 2% low-melt agarose, washed with PBS, and then fixed in 1% osmium tetroxide in cacodylate buffer for 1h. After that, the bacterial cells were post-fixed with osmium tetroxide, dehydrated in gradient-concentration ethanol and embedded subsequently in Epon 812. Specimens were sectioned with a diamond knife into 95nm and they were finally analyzed using a transmission electron microscope S-3400N.

### Metabolite Extraction

Biofilm samples were isolated from 50ml of culture solution, and the supernatants in the culture medium were spun down at 3500rpm for 10min to harvest the planktonic cells. Firstly, each biofilm was clean with PBS for 3 times, and added 2ml the ice-cold methanol to mix by homogenizer for 2min, all procedures were repeated for 3 times, but the samples were frozen and thawed within liquid nitrogen during the process. Secondly the samples were centrifuged under12000 rpm at 4° for 10min. The supernatants were mixed with 800ul of ice-cold acetonitrile for 15min before lyophilization. The dried samples were stored in −80° for GC-MS and LC-MS assay.

### Sample Derivation for GC/MS based Metabolome Assay

The lyophilized samples were resuspended in 40 ul of methoxyamine hydrochloride in pyridine (20 mg/ml) and completely mixed for 90 min at 30°C. N-Methyl-N-(trimethylsilyl) trifluoroacetamide (MSTFA) (90ul) was added to each sample, then mixed further for 30 min at 37°C, and they were undergoing vortex before incubation. The derivative samples were centrifuged at 12000rpm for 10min and then transferred into a 2 ml GC vial and tightly capped for metabolome assay.

### Data acquisition and analysis of GC/MS based metabolome assay

Metabolome assay was carried out on an Agilent 7890-5975C GC/MS system using helium as the carrier gas at a constant flow. A total of 1ul of sample was injected into a 50 m x 250 μm x 0.25 μm DB5-MS column using split less injection. The injector temperature was at 275 °C. Front-Pressure, average speed, flow rate was set at 22.468 psi, 34.908 cm/sec and 1.5 mL/min, respectively. The GC/MS data was firstly transformed to a digital data using XCMS online database and then identified the metabolites by combining AMDIS software and associated database, which based on their retention index (RI) values along with the corresponding mass spectra. The identified metabolites were listed in a data-matrix file that was incorporated with metabolite ID, sample ID and the abundance of each metabolites, it was used for further pattern-recognition and statistical analyses.

### Data acquisition and analysis of LC/MS based metabolome assay

Lyophilized samples were resuspended in 100ul water and was injected to a UFLC-MS/MS system (AB SCIEX API 3200 TQ MS System; UFLC-SHIMADZU, 20AD) equipped with an ESI source in both negative and positive ion-modes with an electrospray ionization voltage of 5500 V for positive mode and 4500 V for negative mode, nebulizer gas (air), turbo gas (air), curtain gas (nitrogen) setting at 50, 50 and 25 psi respectively. Chromatographic separation was carried out on a Waters XSelect HSS T3 column (2.1*100mm, 3.5um) with a gradient program as follows: 5-95% Acetonitrile (Mobile Phase-B) for 50 min at a flow rate of 0.4 ml/min. The column temperature was maintained at 40°C, Mobile-Phase-A: 0.1% formic acid in water and Mobile Phase-B: 0.1% formic acid in acetonitrile. AB SCIEX Analyst software was used for data acquisition and preprocessing. The targeted MRM method was developed based on the acquired reference compounds, which assisted in targeted profiling the known compounds of interest from the known metabolic pathways.

### Statistical Analysis

The data matrix was transferred to MetaboAnalyst 3.0 version online to engage in multivariate statistical analysis including unsupervised principal component analysis (PCA), partial least squares discriminant analysis (PLS-DA) and heat-map overview. The loading plot (VIP plot) was used to discover and identify the differential metabolites, as well as the score plot was adopted to facilitate group classification. This brief method was referred to the publication ^3^

## Results and discussion

### Morphological Observation Verified Successful Development of Biofilm Model

Since notable morphological shifts of biofilm, pellicle and planktonic population were revealed in previous study (Hung et al., 2013). We were first to confirm whether biofilm pellicle model was successful or not in our study (Figure 1B), we employed the cytological method including Crystal Violet Staining, TEM (transmission electron microscope) and SEM (scanning electron microscope), to investigate the intricate-structure phenotype of biofilm formation, as it was expectedly found that pellicle biofilm have an organized and complex extracellular matrix-structure that encased individual, and spatially segregated bacterial-subpopulations (Figures 1E and F), but such phenotype disappeared in the relevant planktonic-subpopulation (Figures 1C and D). This result was completely consistent with previous study (Hung et al., 2013). The imaging assay surely confirmed the biofilm pellicle model was successfully established in this study. In addition, staining technologies such as crystal violet and congo red were used to quantitatively assess the biofilm formation that was also significantly different from the planktonic population (Figure S1). Those phenotypic observations may suggest us there is modified metabolism in two conditions.

### Distinctly metabolic modifications were observed in both biofilm formation and associated planktonic-population

Metabolomics is an systems-biology driven omics method that was designed to global profiling all the small-molecule metabolites, whose level changes are capable of capturing and snapshotting different biological events and processes in cells with a diverse biological contexts (Hobley et al., 2014;Kendall and Sperandio, 2014). To investigate reprogrammed-metabolism underlying biofilm formation, we explored one combinational strategy with untargeted and targeted metabolomics for deciphering the distinct metabolism that was critical for biofilm formation. Metabolome assay revealed that distinctly metabolic reprogramming during biofilm formation, which is significantly different from that in planktonic population (Figure 2). From the score plot resulted from PLS-DA analysis of metabolome data harvested by LC/MS/MS system (Figure 2A), we can see principal component 1 (PC-1 accounted for 31.3% of the variance, with PC-2 explaining 34%. The score plot by GC/MS based metabolome assay (Figure 2B) indicated, principal component 1 (PC1) explained 58.9% of variance, with PC2 of 18.4%, which exhibit the greatest difference predominantly along the PC-1 analysis. Global metabolome result was also illustrated in Figure 2C. Those results revealed remarkable metabolic-differentiation between biofilm formation and planktonic-subpopulation, while bacterial cells were switched to form biofilm, they could secret the extracellular polymeric substance (EPS), which can be used to protect biofilm to defense environment insults and difficult to dissolution. Although our TEM images only recognize the denser layer of fibers (Hung et al., 2013) (highlighted in red in Figure 1E and F) surrounding bacteria in biofilm pellicle mode, many other components (eDNA, proteins and polysaccharides, *etc)* present in EPS structure (Hung and Henderson, 2009;Hobley et al., 2014;Kendall and Sperandio, 2014) which were disappeared in planktonic population. We deduced that microbes are capable of sensing the density of microenvironment as well as releasing the signal to change the gene expression and further yielding the influence upon bacterial metabolism. Consequently, reprogrammed-metabolism would have feedback to the modification with morphological features of the biofilm. In addition, the data also suggested that there was a mechanistic association between biofilm metabolism and its’ morphological feature.

**Figure 2.**
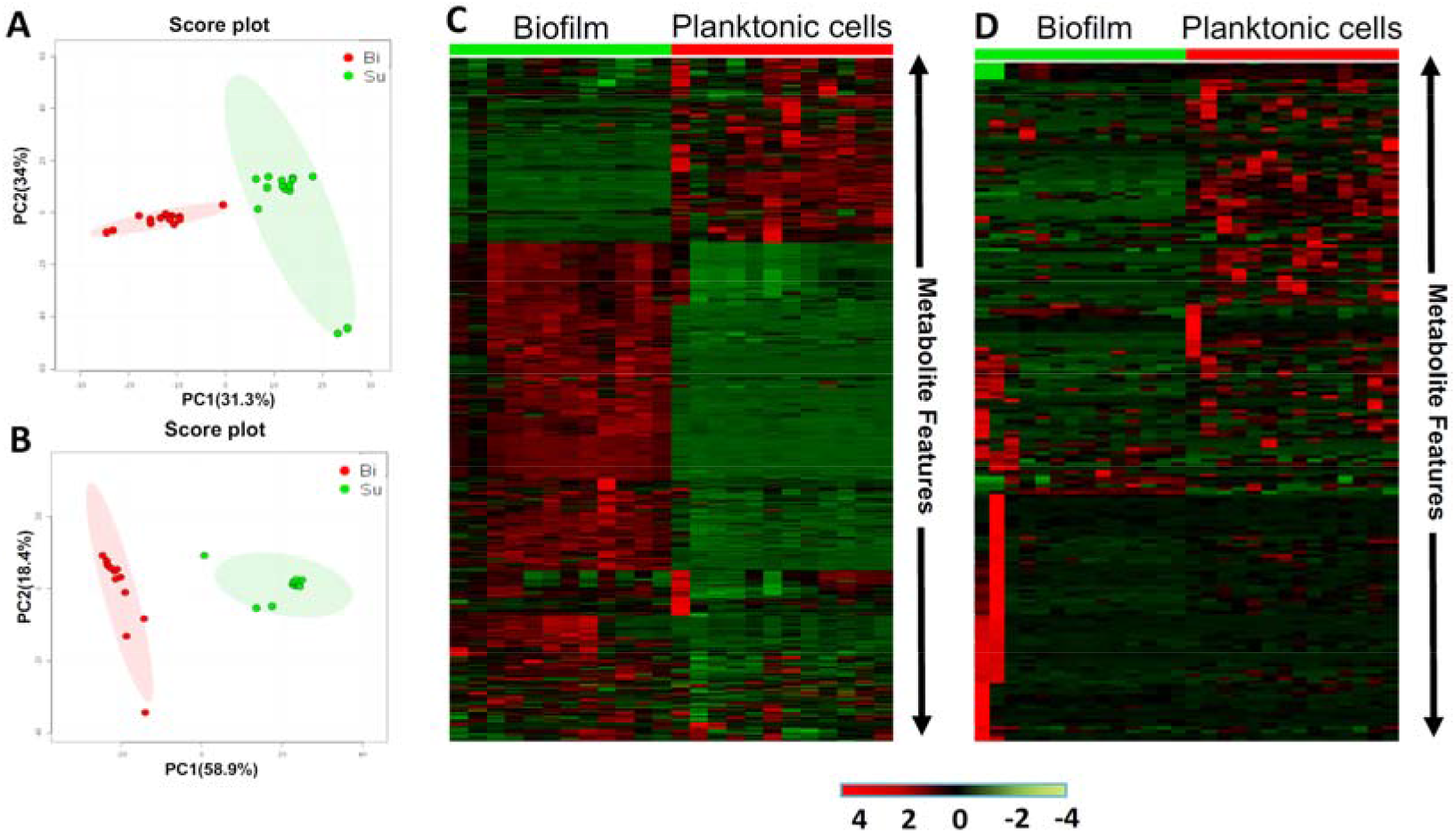
Metabolome assay revealed distinctly modified metabolism between biofilm and planktonic population. (A) Score plot resulted from unsupervised PCA analysis of LC/MS based metabolomics data. (B) Score plot resulted from unsupervised PCA analysis of GC/MS based metabolomics data. (C) Heatmap overviews of global metabolome data with upper one by LC/MS and bottom one by GC/MS. SU: planktonic cells; Bi: Biofilm

### Metabolic reprogramming triggered biofilm formation

Metabolic reprogramming is supposed to be the key driven-factor for biofilm formation even though such organism was structured by the involvements of multiple-layers molecules. We firstly demonstrated obvious modifications in both of small-molecule metabolism and morphological feature between biofilm and planktonic cells, however the profoundly metabolic mechanism remains largely unexplored. Here, we further combined the open-source databases with local database (unpublished) to identify the pivotal metabolites and associated metabolic pathways that have capacity to drive the biofilm formation while compared to planktonic cells. Unexpectedly, we successfully identified 38 differential metabolites, including amino acids, carbohydrates, organic acids, glycerol-derived, polyamine and uridine between biofilm pellicle and planktonic cell (see Table S1 and Figure S1). It was an amazing discovery as glycerol-derived metabolites (2-Phosphoglyceric acid, Glycerol 3-phosphate and D-Glyceraldehyde 3-phosphate) were significantly up-regulated during biofilm formation while the glycerol was decreased considerably. Similarly, we also found that the levels of carbohydrates were enhanced clearly except that the levels of D-Maltose and D-Glucose were decreased in biofilm formation in relevant to that in planktonic cells. The level changes of other differential metabolites in both modes were showed in Table 1 and Figure 3D. Next, to home those differential metabolites to their metabolic pathways, we subjected them to metaboanalyst online to annotate associated metabolic pathways (Figure 4). What metabolism we expected that were mostly reprogrammed that involved glycolipid metabolism, amino acid metabolism and carbohydrate metabolism, which were observed to markedly drive biofilm formation as most of differential metabolites were up-regulated accordingly (highlighted in blue in Figure 4).

**Figure 3.**
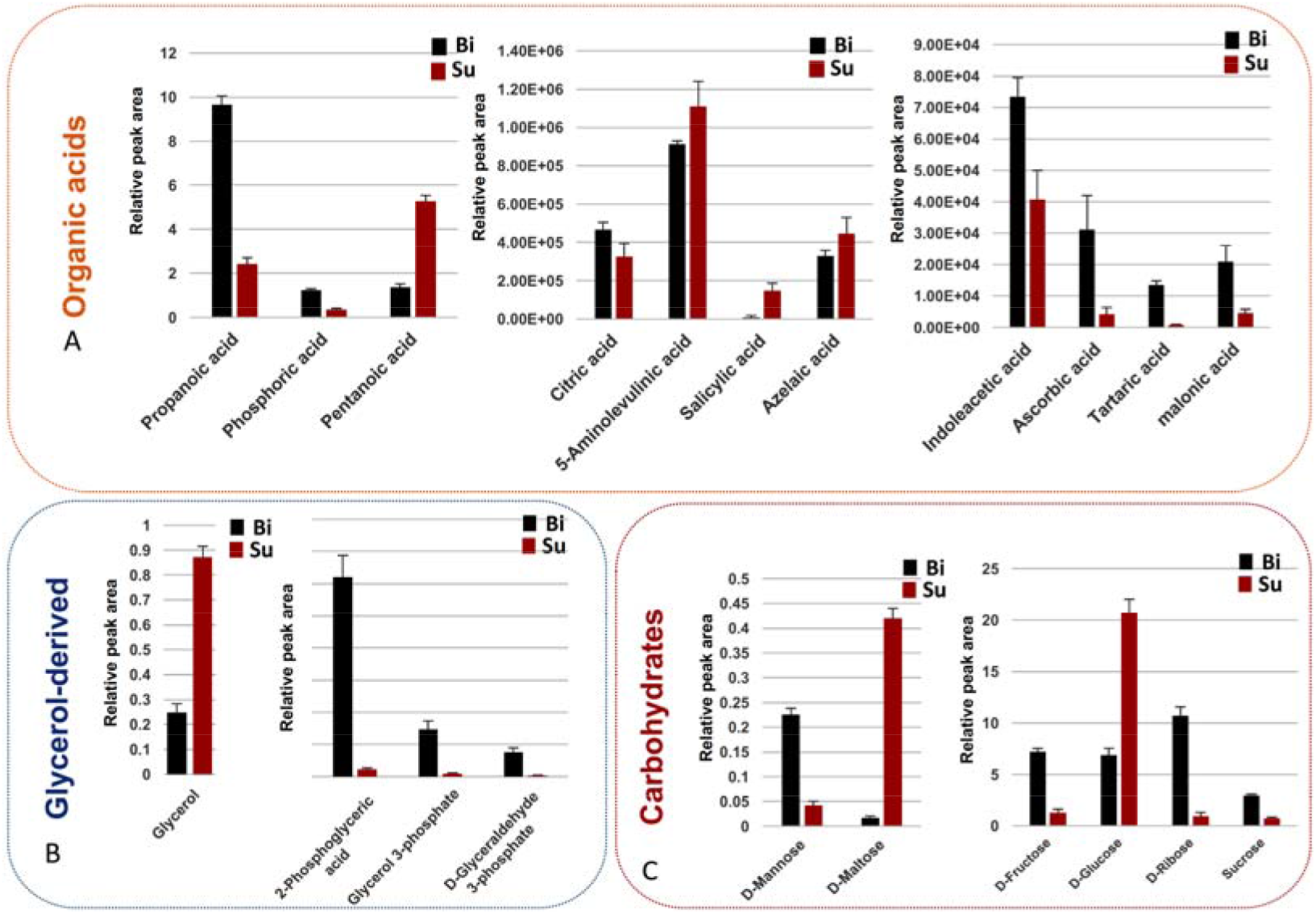
Identifies and their level changes of differential metabolites underlying metabolic reprogramming during biofilm formation while compared to planktonic population. Those differential metabolites mainly include organic acids (A), glycerol-derived molecules (B), and carbohydrates(C). SU: planktonic cells; Bi: Biofilm

**Figure 4.**
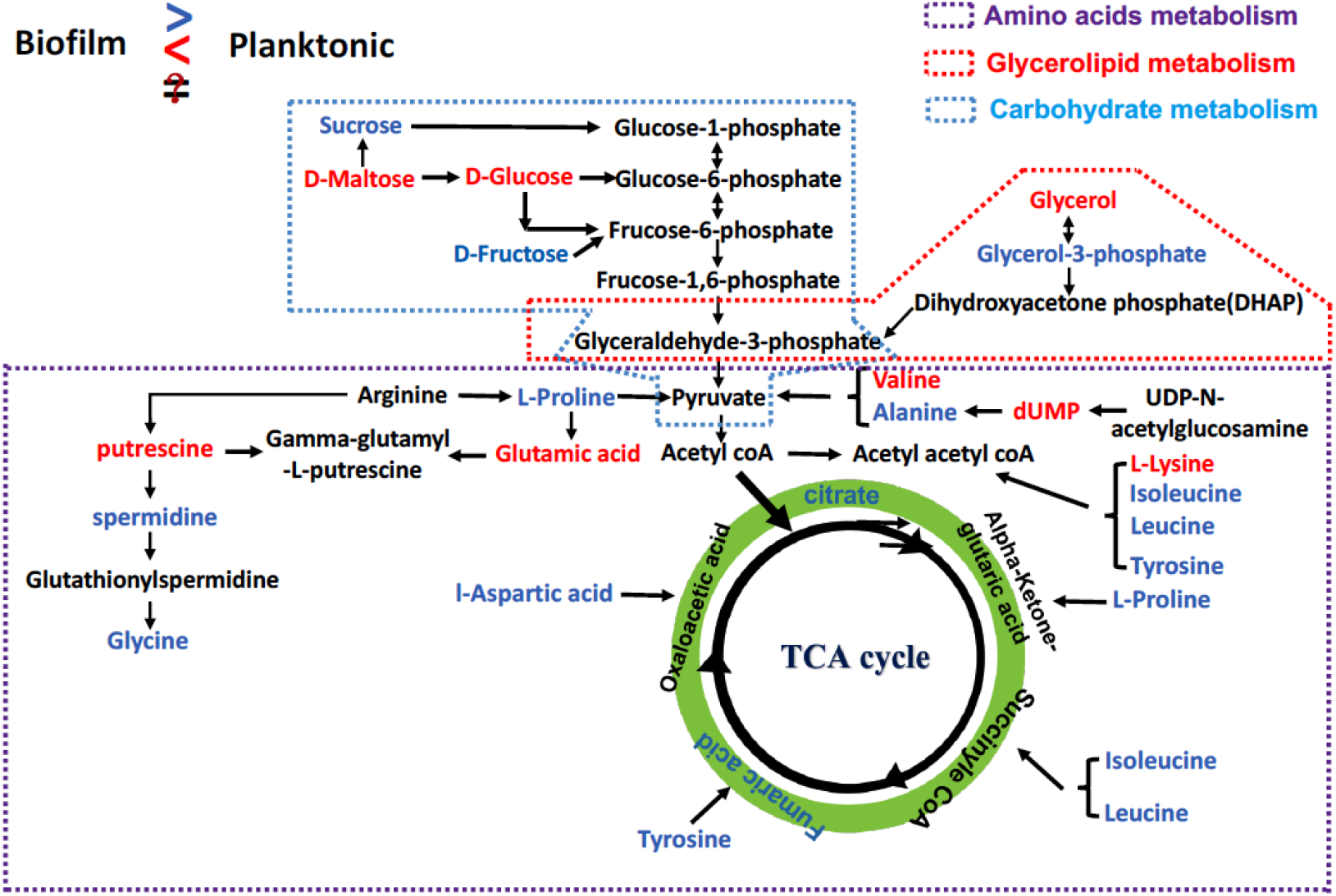
The mostly affected metabolic pathways triggered substantially metabolic reprogramming that permitted biofilm formation induced by UTI89 strain. Amino acid metabolism, glycerolipid and TCA cycle are mostly affected during biofilm formation as differential metabolites in blue were significantly increased in biofilm formation, and blue ones were lowered remarkably.

It was reported that proteins and carbohydrates were the main components of biofilm formation (Costerton et al., 1995). Several amino acids were documented to account for biofilm detachment and maturation (Valle et al., 2008;Kolodkin-Gal et al., 2010;Bernier et al., 2011;Leiman et al., 2013). In addition, the mixture of D-Amino acids would lead to the biofilm desperation with *Bacillus subtili* and further break down the biofilm by YqxM protein which regulated the amyloid fibers anchoring or off-anchoring to the host cells (Kolodkin-Gal et al., 2010). However, our results only found the levels of many L-amino acids rather than D-amino acids were sharply changed (Figure 3C), which might reveal the L-amino acids may be also participated in biofilm formation induced by UTI89 (Leiman et al., 2013) while D-amino acids were catalyzed as dispensable nutrition for bacteria growth (Bayne and Stokes, 1961;Fernandez and Zuniga, 2006). In addition, we also noticed that the level changes of different amino acids were extremely distinct between biofilm and planktonic cells. This unregularly phenomenon supported again that amino acids may specially play their individual roles in biofilm formation(Wong et al., 2015). Metabolic reprogramming of amino acids in this study was confirmed again to mostly trigger biofilm formation.

AI-2 is a typically quorum-sensing (QS) molecule, it can regulate physiological process of biofilm formation as the density of bacterial cells changed (Kendall and Sperandio, 2014). The lsr protein is essential for importing and processing AI-2, which was noticed to be inhibited by high concentration of Glycerol-3-phosphate (G-3-P)(Willias et al., 2014). We speculated that biofilm formation preferred to consume glycerol to biosynthesize G-3-P as it could regulate the expression of AI-2 to avoid biofilm dispersal. This was agreeable with our result that the level of glycerol during biofilm formation was decreased remarkably while the production of G-3-P was enhanced substantially (Figure 3B). Moreover, the most carbohydrates were incorporated into FimH adhesion (mannose) to increase the adhesive capability of biofilm (Schwartz et al., 2013) or participate in the EPS formation. So bacteria would seek other nutrients to maintain their survival. The G-3-P was the key precursor of glyceraldehyde-3-phosphate, it was a key factor that connected the carbohydrate metabolism to glycerolipid metabolism, as well as was a candidate for providing the microbes with the necessary energy. Thus, our finding convinced that more G-3-P has been synthesized, via consuming more glycerol to maintain the survival of bacterial cells during biofilm formation.

Interestingly, our data showed that putrescine was expressed differentially between two modes of UTI89. This result was well supported by previous studies that putrescine plays a crucial role in modulating the biofilm formation triggered by some bacteria(Burrell et al., 2010;Wortham et al., 2010;Goytia et al., 2013;Hobley et al., 2014). Another polyamine spermidine was observed to have a remarkable shift in biofilm formation, which was not expected since spermidine was verified only to inhibit the growth of *E. coli* at PH 7, but not link to the biofilm formation(Nesse et al., 2015). In near future, we shall figure out to study further the mechanisms of this metabolite modulating biofilm formation.

Bacterial cells in biofilm mode have extraordinary survival ability (Chauret, 2011) to resist the environmental stress largely via the protective EPS. Amino acid metabolism, carbohydras metabolism and glycolipid metabolism in bacteria were not only responsible for the biosynthesis and storage of accessible energy but also were utilized for synthesizing the secreted components such as amino acids, sugars, lipids, uridines and organic acids, which are essential for EPS production during biofilm formation (Larsen and Engelsen, 2015). Those metabolic pathways were highlighted in this study to have significantly metabolic reprogramming that triggered the biofilm formation from metabolic perspective (Figure 4). Those findings indicated that biofilm formation is ought to yield a protective feature, by funneling metabolites into EPS synthesis, conveying the stronger protection against stress conditions via dominating the metabolic reprogramming. Biofilm network is supposed to build a anaerobic microenvironment for bacteria embedded in biofilm pellicle (Hamilton, 1987), where bacterial cells were structured in a stable space. Thus, bacterial biofilm could secure more resources in order to survive well while compared to planktonic cells.

## Conclusion

In summary, this study was first to combine metabolomics with cytology methods together to better understand the biofilm formation in UTI89 strain. Our data has revealed that obviously metabolic reprogramming triggered biofilm formation while compared to planktonic population as we found lots of small-molecule metabolites are critically essential for biofilm formation and dissociation induced by UTI89. Those differential metabolites and the associated metabolic pathways can be regarded as novel targets for the development of biofilm based treatments and antibiotic discovery against the UTI89 causing infection. More importantly, such effort could provide a novel insight into better understanding of the biofilm formation caused by different bacterial strains at metabolic level.

## Acknowledgements

This work was supported by the National Natural Science Foundation of China Grants (No. 81274175 and c010201), National Key R&D Program of China (No. 2017YFC1308600 and 2017YFC1308605), and the Startup Funding for Specialized Professorship Provided by Shanghai Jiao Tong University (No. WF220441502).

## Author Contributions

H.T.L. and Y.M.Q. designed and conceived the experiments; Y.M.Q., X.W., T.B.G., H.G. and H.T.L. collected and analyzed the data; H.T.L. and Y.M.Q. wrote the manuscript. All authors reviewed the manuscript.

