## Supporting Table S1, Figure S2 and Figure S3 for "Mass Spectrometry based Metabolomics Deciphered Metabolic Reprogramming That Was Required for Biofilm formation in Uropathogenic *Escherichia coli*"

**Supporting Table S1 and Figures S1-2**

**Mass Spectrometry based Metabolomics Deciphered Metabolic Reprogramming That Was Required for Biofilm formation in Uropathogenic *Escherichia coli***

Haitao Lu^1*#^, Yumei Que^2#^, Xia Wu^2^, Tianbing Guan^2^, Hao Guo ^2^

^1^Key Laboratory of Systems Biomedicine (Ministry of Education), Shanghai Center for Systems Biomedicine, Shanghai Jiao Tong University, Shanghai 200240, China

^2^School of Pharmaceutical Sciences, Chongqing University, Chongqing 401331, China

^#^The authors are equally contributed to this article

*Corresponding author.

Haitao Lu Ph.D. Professor, Key Laboratory of Systems Biomedicine (Ministry of Education), Shanghai Center for Systems Biomedicine, Shanghai Jiao Tong University, Shanghai 200240, China

**Table S1.** Identifies of differential metabolites characterized by both of LC/MS and GC/MS based metabolomics assay of biofilm and plantonic cells.

| Classification of Metabolites | Match | | HMDB | | PubChem |
| --- | --- | --- | --- | --- | --- |
| Carbohydrates | D-Fructose | HMDB00660 | | 439709 | |
|  | D-Glucose | HMDB00122 | | 5793 | |
|  | D-Mannose | HMDB00169 | | 18950 | |
|  | D-Ribose | HMDB00283 | | 5779 | |
|  | Sucrose | HMDB00258 | | 5988 | |
|  | D-Maltose | HMDB00163 | | 10991489 | |
| Amino Acids | Glycine | HMDB00123 | | 750 | |
|  | L-Valine | HMDB00883 | | 6287 | |
|  | L-Isoleucine | HMDB00172 | | 6306 | |
|  | L-Proline | HMDB00162 | | 145742 | |
|  | L-Lysine | HMDB00182 | | 5962 | |
|  | L-Alanine | HMDB00161 | | 5950 | |
|  | L-Leucine | HMDB00687 | | 6106 | |
|  | L-Aspartic acid | HMDB00191 | | 5960 | |
|  | L-Tyrosine | HMDB00158 | | 6057 | |
|  | L-Methionine | HMDB00696 | | 6137 | |
|  | L-Glutamic acid | HMDB32478 | | 32881 | |
|  | N-Acetylglutamic acid | HMDB01138 | | 185 | |
|  | L-Phenylalanine | HMDB00159 | | 6140 | |
| Uridine | dUMP | HMDB00071 | | 65058 | |
|  | 2'-Deoxyguanosine 5'-monophosphate | HMDB02894 | | 65040 | |
| Polyamine | Putrescine | HMDB01414 | | 1045 | |
|  | Spermidine | HMDB01257 | | 1102 | |
| Glycerol-derived | Glycerol | HMDB00131 | | 753 | |
|  | 2-Phosphoglyceric acid | HMDB00362 | | 59 | |
|  | Glycerol 3-phosphate | HMDB35909 | | 435280 | |
|  | D-Glyceraldehyde 3-phosphate | HMDB01112 | | 3418 | |
| Organic acids | Citric acid | HMDB00094 | | 311 | |
|  | 5-Aminolevulinic acid | HMDB01149 | | 137 | |
|  | Salicylic acid | HMDB01895 | | 338 | |
|  | Azelaic acid | HMDB00784 | | 2266 | |
|  | Salicylic acid | HMDB01895 | | 338 | |
|  | Azelaic acid | HMDB00784 | | 2266 | |
|  | Indoleacetic acid | HMDB00197 | | 802 | |
|  | Tartaric acid | HMDB14433 | | 443958 | |
|  | Malonic acid | HMDB00691 | | 867 | |
|  | Propionic acid | HMDB00237 | | 1032 | |
|  | Phosphoric acid | HMDB02142 | | 1004 | |

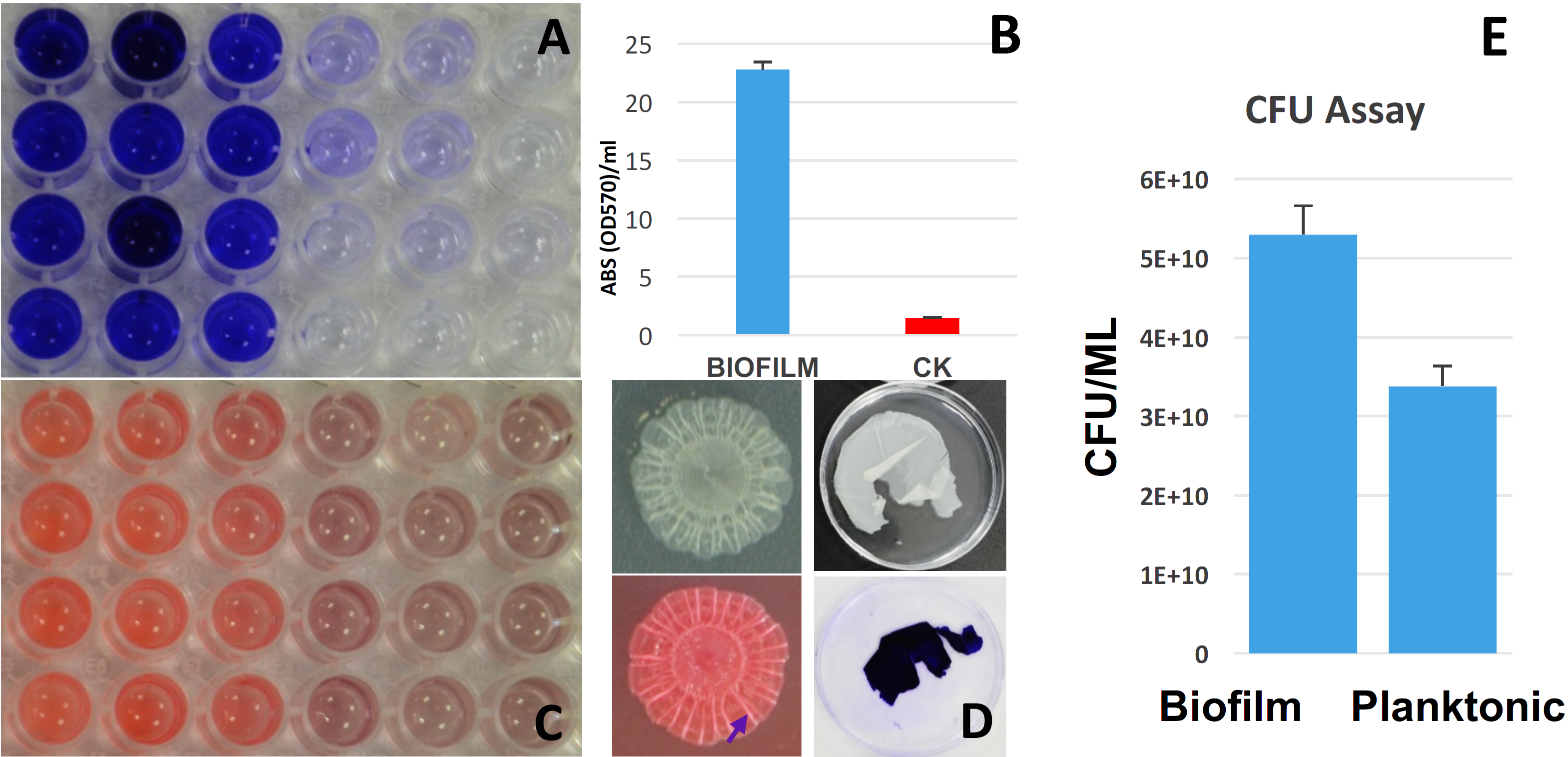

**Figure S1.** Staining assays were employed to phenotype the biofilm formation. Crystal violet (A). Absolute value (OD570nm) of crystal violet in (A) was quantitatively measured (B). Congo red (C). Biofilm based on agar culture, and biofilm stained with congo red in left two panels; biofilm based LB culture and biofilm stained with crystal violet based LB culture in right two panels (D). Colony-Forming Units were measured in both biofilm and planktonic cells (E).

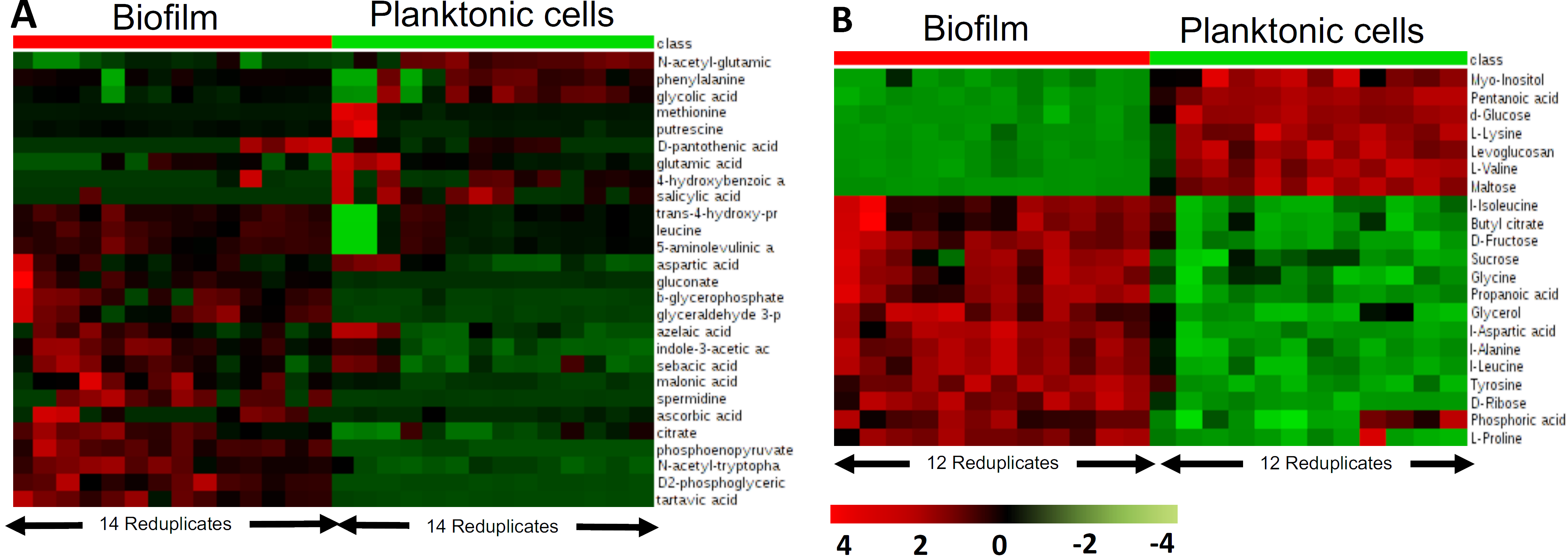

**Figure S2.** Metabolome assay revealed significantly metabolic reprogramming during biofilm formation. (A) Heatmap overview of differential metabolites identified by LC/MS based targeted metabolome asaay**.** (B) Heatmap overview of differential metabolites identified by GC/MS based untargeted metabolome asaay.
